# Genome assemblies of climatically distinct *Aegilops mutica* reveal extensive structural variation

**DOI:** 10.64898/2026.09.28.753720

**Authors:** Ashley L. Tuffin, Georgios Banos, Benjamin Kilian, Wayne Powell, Alex D. Twyford, Matthew Hartfield, Rajiv Sharma

## Abstract

Bread wheat harbours low genetic diversity due to bottlenecks during speciation, domestication, and selection; wild relatives therefore provide critical reservoir for climate-resilient breeding. *Aegilops mutica*, one of the few self-incompatible diploids in the *Aegilops–Triticum* species complex, is a putative parent in the homoploid hybridisation event that formed the D-genome clade and is a proven donor of disease resistance, winter hardiness and grain zinc accumulation. *Ae. mutica* is globally endangered, with a reference-quality genome available for only a single Central Anatolian accession. Using HiFi sequencing, we generated genome assemblies from two accessions from opposite extremes of the species’ range: western Türkiye and Armenia. The four haploid assemblies recovered ~93% complete BUSCOs and comprised ~81% repetitive sequence, predominantly LTR retrotransposons (~55%), closely matching the reference. Comparison against the reference identified 145,871–155,535 structural variants per haplotype, including an 8.4 Mb inversion on chromosome 6T. Between 96,860 and 105,472 structural variants segregated between haplotypes within individuals and classic domestication loci did not overlap with any large structural variants. These assemblies and structural variant catalogue extend genomic resources for *Ae. mutica*, providing a basis for targeted introgression into wheat and conservation of a species for which populations in the South Caucasus have severely declined.

**Plain language summary:** Bread wheat has limited genetic diversity due to selection, making it vulnerable to climate change and disease. Crop wild relatives contain useful variation that could be used to breed more resilient crops. *Aegilops mutica* is an endangered wild grass related to wheat that has been used before for disease resistance and other traits. We sequenced two plants originally collected from opposite ends of its range – western Türkiye and Armenia. We found over 100,000 large-scale differences per plant between our new genomes and the reference genome, including one rearrangement over 8 million DNA bases long. Even within one plant, the two copies of its genome contained many differences, as *Aegilops mutica* must always cross-pollinate to reproduce. These findings increase our knowledge of this endangered species, providing wheat breeders with additional resources and conservationists with a baseline of its genetic diversity across its range.

## Introduction

Crop species often face reduced genetic diversity due to domestication, leading to vulnerability to biotic and abiotic stresses (Bohra *et al*., 2022; Dempewolf *et al*., 2017; Brozynska *et al*., 2016; Kishii, 2019). As global agriculture faces the dual pressures of climate volatility and rising food demand, the need to increase the narrow genetic base of elite cultivars has become a primary concern. Bread wheat (*Triticum aestivum* L.) exemplifies this concern, having undergone successive genetic bottlenecks during speciation, domestication, and intensive human selection that significantly depleted its standing variation (Kihara, 1919; Sax, 1921; Sax, 1922; Kilian *et al*., 2014; Levy and Feldman, 2022) (Figure 1a). Crop wild relatives (CWRs) offer a solution to this challenge by providing a rich source of genetic diversity for breeding programs (Redden *et al*., 2015; Kilian *et al*., 2021). Ensuring food security requires the development of resilient, sustainable, and efficient cultivars, and the genetic diversity within crop wild relatives must therefore be characterised and preserved (Castañeda-Álvarez *et al*., 2016). The secondary gene pool (Harlan and de Wet, 1971) of wheat is a prime example of CWRs that have been utilised in breeding programs, however, there remains untapped genetic diversity in wild species that have so far been overlooked (Sharma *et al*., 2021). Mobilising the lost diversity within the secondary and tertiary gene pools is therefore a critical mandate for climate-resilient breeding (Nevo, 2006; Kilian *et al*., 2011; Stalker *et al*., 2021; Eastwood *et al*., 2022).

**Figure 1.**
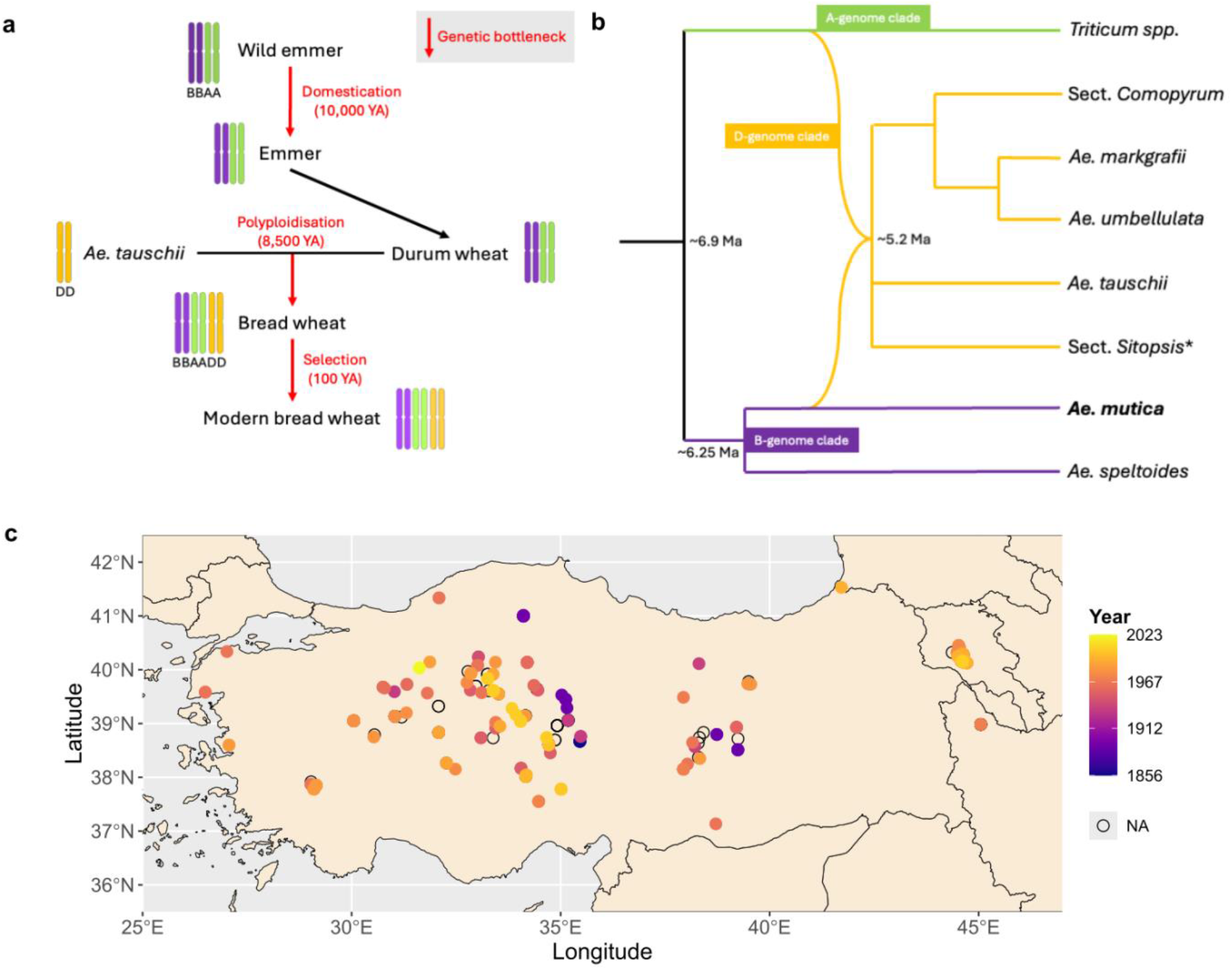
Evolutionary origins and geographic distribution of *Ae. mutica*. a) Illustration of the genetic bottlenecks (red) in bread wheat necessitating the incorporation of wild relative diversity. b) Schematic representation of the Aegilops-Triticum species complex illustrating the pivotal position of *Ae. mutica* and its potential role in the homoploid hybridization event leading to the D-genome clade. *Triticum, Comopyrum*, and *Sitopsis* clades collapsed (asterisk highlights the exclusion of *Ae. speltoides* from section *Sitopsis*). Approximate dates from Glémin *et al*. 2019, phylogeny from Bernhardt *et al*. 2020. c) Genesys and GBIF observations of *Ae. mutica* across the Anatolian peninsula and South Caucasus, coloured by year recorded.

*Aegilops mutica* Boiss. (or *Amblyopyrum muticum* (Boiss.) Eig) is one of the few self-incompatible, outcrossing diploids in the *Aegilops-Triticum* complex and maintains high levels of heterozygosity and structural diversity that remain largely undescribed in cereal genomics (Burgarella *et al*., 2023; Grewal *et al*., 2025). Despite its unique evolutionary position as a putative parent in the homoploid hybridisation event that formed the D-genome clade (Glémin *et al*., 2019; Bernhardt *et al*., 2020) and being sister species to *Aegilops speltoides*, the closest living relative of the wheat B-genome donor (Glémin *et al*., 2019; Huynh *et al*., 2019; Bernhardt *et al*., 2020; Zhang *et al*., 2022) (Figure 1b), *Ae. mutica* remains critically underrepresented in global genebanks and genomic databases (Crop Trust, 2017). This lack of data represents a major gap that obscures our understanding of how standing genomic variation is maintained and distributed across the species’ range.

Historically, *Ae. mutica* had a distribution stretching across the entire Anatolian peninsula and through the South Caucasus to the coast of the Caspian Sea (<u>Figure 1</u>c), where it was collected and observed up until the late 1900s (Kilian *et al*., 2011; van Slageren, 1994; Aminov and Aliyeva, 2012; Eldarov *et al*., 2015; GBIF Secretariat: GBIF Backbone Taxonomy, 2023). Now, it is globally endangered – its range limited to Central Anatolia and 1-2 critically endangered populations in the South Caucasus (Haruntyunyan *et al*., 2010; Brehm, 2017). Prior genomic characterization of *Ae. mutica* has been largely confined to its utility within *Triticum* introgression programs (Iefimenko *et al*., 2018; King *et al*., 2017; King *et al*., 2019; Othmeni *et al*., 2019) and description of SNP diversity from cross-pollinated genebank accessions (Adhikari *et al*., 2023). *Ae. mutica* genetic resources have already proven invaluable for identifying a suite of high-value traits, including resistance to powdery mildew and wheat rusts (Eser, 1998; Liu *et al*., 2016; Fellers *et al*., 2020; Xu *et al*., 2025), enhanced winter hardiness (Iefimenko *et al*., 2015), and superior grain zinc accumulation (Guwela *et al*., 2024). While contig-level and phased chromosome-level assemblies have recently been published for a single accession, T2130012, from Nevşehir Province (Coombes *et al*., 2023; Grewal *et al*., 2025), the broader genomic landscape of the species remains largely unresolved. Although the population structure within Central Anatolia has previously been assessed via simple sequence repeat (SSR) markers (Alsaleh *et al*., 2022), the genomic diversity of the species requires the context of multiple assemblies to fully understand genome-wide variation such as structural variation (SV). The large size of wheat genomes has been a barrier to research due to prohibitive costs associated with sequencing, however, advances in sequencing technologies coupled with innovative studies in low-coverage methods have opened an avenue for exploring non-model species with large genomes (Hu *et al*., 2025).

Recent pangenome studies in *Avena* and *Hordeum* demonstrate that SVs, rather than SNPs, frequently underlie rapid adaptation by creating non-recombining supergenes (Feng *et al*., 2025; Zhang *et al*., 2025). In outcrossing species like *Ae. mutica*, these structural polymorphisms are maintained at high frequencies, potentially harbouring distinct adaptive strategies for the cold steppe versus Mediterranean climates. Large haplotypes, also often formed via inversion SVs, can be correlated with ecotypic adaptation in plants, as seen in *Ambrosia, Gossypium, Helianthus*, and *Mimulus* (Todesco *et al*., 2020; He *et al*., 2021; Battlay *et al*., 2023; Lowry and Willis, 2010). Since the historical range of *Ae. mutica* spans several distinct climatic zones, a single reference assembly is unlikely to capture the structural diversity of the species. We used bioclimatic clustering of the known species range to guide sampling design; accessions that are climatically and geographically distinct from the reference are more likely to harbour unrepresented structural variation if they underpin local adaptation.

Here, we move beyond a single reference genome by generating four new partially phased assemblies from two individuals of accessions originally collected in distinct, unquantified climates. Our goal was to compare these new assemblies to the reference genome (Grewal *et al*., 2025) and identify structural variants across assemblies, which represent candidate loci for future assisted-breeding programs and could underlie local adaptation. We hypothesised that accessions originating from climatically distinct regions would harbour substantial structural variation not represented in the current reference genome. To do this, we used bioclimatic clustering of georeferenced occurrence records to select two accessions from opposite range edges, generated partially phased assemblies from low-coverage (~9× per haplotype) PacBio HiFi data, and scaffolded them to the T2130012 reference. We then called structural variants with SyRI – both against the reference and between the two haplotypes of each individual – annotated repeat content with EarlGrey, and tested SV-overlapping genes for GO enrichment and for presence of classic wheat domestication loci.

## Materials and Methods

### Bioclimatic analysis

We used bioclimatic clustering to prioritise accessions most likely to contain unrepresented structural variation. We compiled the locations of all Ae. mutica observations and collections recorded on GBIF (GBIF.org, 2023) and Genesys (Crop Trust, 2017) with observations from the literature (Alsaleh *et al*., 2022). In some cases where coordinates were not provided but the site location was described, coordinates were estimated. Using the WorldClim dataset (Fick and Hijmans, 2017), we retrieved 19 bioclimatic variables for each of the 295 geolocated observations or collections. We performed a principal component analysis on the scaled bioclimatic variables and applied k-means clustering to the resulting principal component scores. We set K = 3, the value at which cluster membership corresponded most closely to the major Köppen-Geiger classifications across the species range. We note that this clustering is used descriptively to structure sampling across the climatic range of the species and is not intended as a statistical test of population or ecotypic differentiation. Locations and cluster assignments were plotted in R using terra (Hijmans *et al*., 2026) and ggplot2 (Wickham, 2016).

### Plant materials

As *Aegilops mutica* is an obligate outcrossing species, genome assemblies were generated from single individuals sampled from each accession; throughout the manuscript we distinguish between accession (germplasm unit) and individual genome where appropriate. Both individuals chosen for genome sequencing are from seed bank collections. Sample T05_6D (or T05), from accession PI 560121/84TK157-0003, was provided by The Western Regional Plant Introduction Station of the United States Department of Agriculture’s Agricultural Research Service (USA022). The accession was originally collected in 1984 from the Çürüksu valley near Pamukkale/Denizli, Türkiye. Sample T19_2C (or T19), from accession AE 912/AE 367 B, was provided by the Leibniz Institute of Plant Genetics and Crop Plant Research (DEU146) and was originally collected in Armenia by Dr. Gandiljan around 1977.

We pre-germinated seeds from both accessions on damp filter paper in petri dishes within a glasshouse. After one week, seeds had germinated and were ready to be individually planted. One month later, we dark-treated the plants for 24 hours as advised by the LeafGo methodology (Driguez, Bougouffa, *et al*., 2021) and collected whole leaf tissue samples before flash freezing them in liquid nitrogen and storing at −80°C.

### DNA extraction and sequencing

To get pure, high quantity, high molecular weight DNA, we used two different extraction methods. For sample T05_6D, we used the Monarch HMW DNA Extraction Kit following the manufacturer’s instructions, however, we isolated DNA using cold centrifugation (Nishii *et al*., 2023) from the isopropanol and NaCl solution that was washed over the glass beads rather than proceeding to the final elution step. For sample T19_2C, we followed the LeafGo protocol utilising the Qiagen Genomics-Tip kit (Driguez, Carty, *et al*., 2021).

We measured concentration and purity using a Qubit and DeNovix spectrometer, respectively. T05_6D was measured at 29.8 ng/μL (7450 ng, total), with 1.79 260/280 and 1.09 260/230. T19_2C was measured at 43.8 ng/μL (10950 ng, total) and 1.94 260/280 and 1.98 260/230. Additional quality control of samples was conducted by Edinburgh Genomics using Qubit and Tapestation. T05_6D and T19_2C were confirmed as having no RNA contamination and DNA Integrity Numbers (DIN) of 8.8 and 9.8, respectively. Both samples had Tapestation peaks measured at near or over 60 kbp and were sheared to roughly 15 kbp before sequencing libraries were prepared and circular consensus sequencing (CCS) was carried out on a PacBio Revio. Over 5 million HiFi reads were generated for each sample, ranging from 80 to 87 Gb. Given an estimated genome size of 4.6 Gb, this corresponds to approximately 9× coverage per haplotype. This depth was chosen deliberately: low-coverage long-read sequencing substantially reduces the per-genome cost of surveying structural variation in large Triticeae genomes, and structural variants can be detected in wheat from HiFi reads with high precision at coverage as low as 0.33× and varied but plateauing recall at 6.60× coverage (Zhang *et al*., 2024).

### Genome assembly

Using the Crop Diversity HPC (Percival-Alwyn *et al*., 2025), we produced two partially-phased assemblies from HiFi CCS fastq reads using hifiasm (v0.24.0-r702) with “-s 0.15 --hg-size 4607m” parameters. We determined the optimal similarity threshold (Cheng *et al*., 2021) of 0.15 through comparisons of BUSCO (v5.8.3, viridiplantae_odb12 database) results for various values of “-s” (Figure S1). Using RagTag (v2.1.0), we then scaffolded the two hifiasm assemblies to the reference assembly, Ae.mutica_hap1_with_organelles (GCA_964657205.1, haplotype 1 of accession T2130012), excluding unplaced scaffolds and inferring gap sizes, then used assembly_stats for quality control and summary statistics (Cheng *et al*., 2021; Alonge *et al*., 2022; Manni *et al*., 2021; Trizna, 2020).

### Structural variant detection

To detect and visualise structural rearrangements and synteny within and between samples, we used SyRI (v1.6.3) and plotsr (v1.1.1) (Goel *et al*., 2019; Goel and Schneeberger, 2022). Chromosomes from both haplotypes of T2130012 were extracted using the subseq function of seqtk (v1.4-r122), and all assemblies were soft masked using tantan (v51) (Frith, 2011). We used MUMmer4 (v4.0.1) to align, filter, and convert outputs to acceptable inputs for SyRI (Marçais *et al*., 2018). Both haplotypes of an individual were compared to each other and to the reference genome, with stated sizes relative to the reference genome or, when between two haplotypes, the haplotype labelled “1”. These were produced using NUCmer with parameters “-l 100 -c 500 -b 500” to reduce hits from repetitive or transposable elements then filtered with delta-filter “-m -i 90 -l 500” to remove low-identity, small alignments before being converted to a tabular format using show-coords “-THrd”. Results from SyRI (default parameters) were plotted using plotsr using various parameters, “-itx” for inter-chromosomal plotting, “-R” for cleaner visualisation, and varying “-s” for minimum variant size to be plotted. We used bcftools (v1.3.1) to extract information on count values of structural rearrangement type and location, then plotted the data using ggplot2 (Danecek *et al*., 2021; Wickham, 2016). To further investigate the effect of structural rearrangements, we used BLAST to locate wheat domestication genes in haplotype 1 of T2130012 to cross-reference the locations with any structural variants in the new assemblies.

### Transposable element detection and GO enrichment analysis

We used EarlGrey (version 4.4.4) (Baril *et al*., 2024a; Baril *et al*., 2024b) to detect repetitive elements such as transposable elements (TEs), utilising the RepeatMasker search term “-r viridiplantae” and a TE library created according to Sun *et al*., 2025 that combined the CLARI-TE database (Daron *et al*., 2014; Daron, 2014) and TREP (the TRansposable Elements Platform) (Wicker *et al*., 2002; Schlagenhauf and Wicker, 2016), with clustering to reduce redundancy “-c” and putative spurious TE annotations removed “-m”.

To carry out gene ontology (GO) enrichment analysis, we first generated a proteome from the T2130012 reference genome assembly and gff3 using AGAT (Dainat *et al*., 2026). We used eggNOG mapper (v2.1.13; Huerta-Cepas *et al*., 2019; Cantalapiedra *et al*., 2021) with DIAMOND (v2.0.15; Buchfink *et al*., 2021) to determine the associated GO terms of reference genes – all reference genes with at least one GO term were used as the background or “gene universe” for enrichment analysis (26,660 of 105,854). GO terms that feature animal-specific language, such as “blood circulation” and “synapse”, are artifacts of annotation transfer and were disregarded. Using bedtools intersect, a reference gene was associated with an SV if the gene annotation overlapped with a large SV by at least one base pair. SVs were called by SyRI relative to the reference and included deletions over 50 bp excluding SNP/short indels. Finally, topGO’s Fisher’s exact test with the weight01 algorithm (Alexa and Rahnenführer, 2009) was used to measure GO enrichment within SVs (nodeSize = 10, Benjamini–Hochberg FDR < 0.1).

## Results

### Bioclimate informed sampling

Bioclimatic clustering of occurrence data identified distinct climatic groupings that we used to guide accession selection. PCA analysis of bioclimatic variables at observation sites from across the species range showed meaningful clustering when *K = 3* (Figure 2a). The clusters largely correspond to different Köppen-Geiger climate classifications (Figure 2b; Beck *et al*., 2018): cluster one consists of cold, semi-arid steppe (BSk), veering into humid continental climate with dry, cool or warm summers (Dsb/Dsa); cluster two is a temperate climate of Mediterranean hot summers in the South-West and humid subtropical in the North (Csa/Cfa); and cluster three, like cluster one, is also a cold, semi-arid steppe (BSk) that veers into a humid continental climate but instead has a mild summer and is wet year-round (Dfb). Clusters also separated spatially: cluster one represents the observations from the interior of Central Anatolia, cluster two mostly aligns with the observations from the exterior of the Anatolian Peninsula, and the population in Armenia corresponds to cluster three (Figure 2c). The reference accession, T2130012, was collected in Nevşehir Province and falls within cluster one. Two accessions collected from clusters two and three were chosen to capture the range edge of the species: T05, from the Çürüksu valley near Denizli, representing cluster two; and T19, from Armenia, representing cluster three.

**Figure 2.**
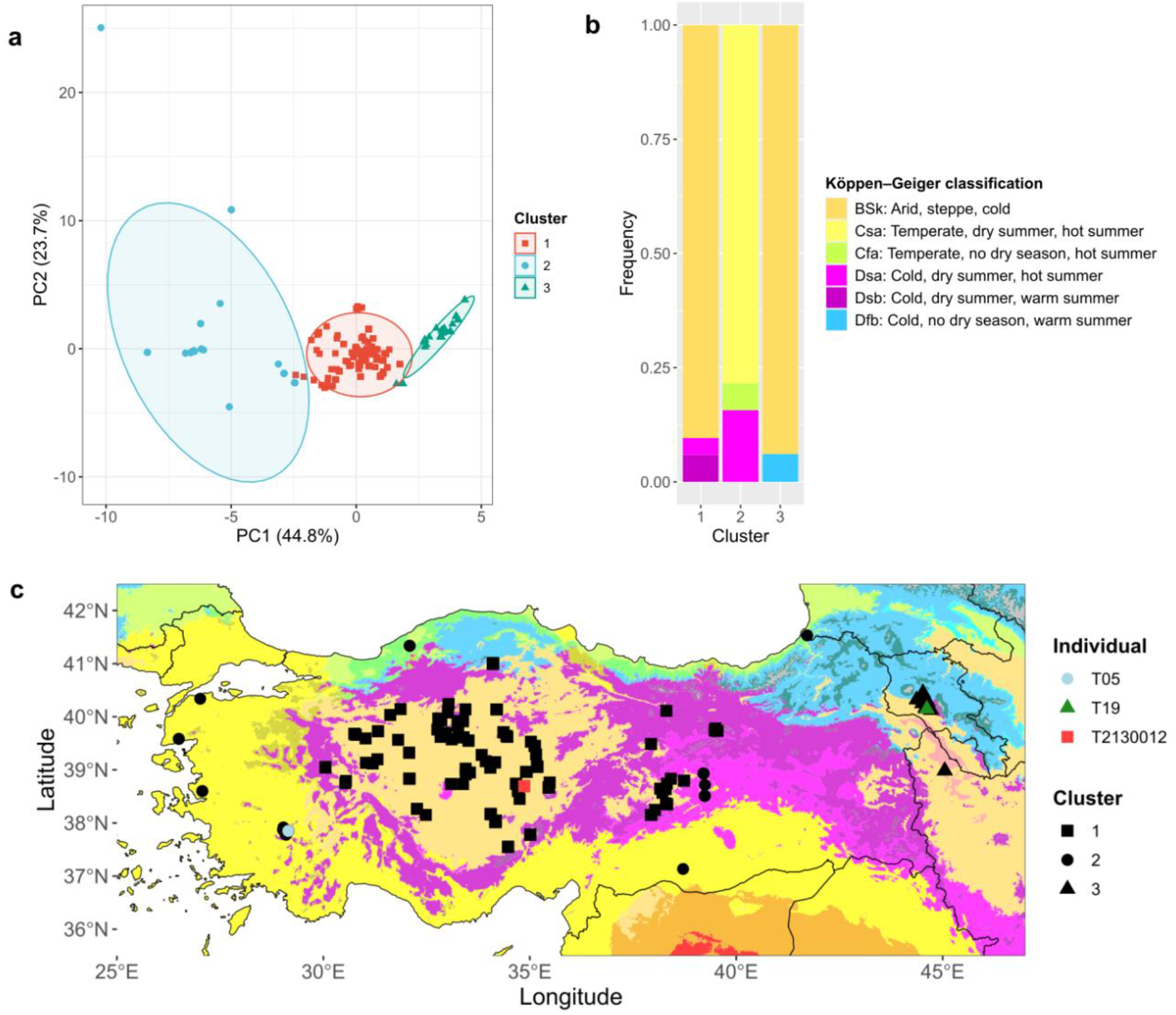
Bioclimatic clustering of *Ae. mutica* accessions. a) Principal Component Analysis of bioclimatic variables with K = 3, identifying three distinct bioclimatic clusters. Variables strongly contributing to PC1 were related to mean and minimum temperature and precipitation in the coldest month/quarter and to PC2 (23.7%) were precipitation in the driest and hottest month/quarter. b) Frequency of Köppen-Geiger climate classifications within each cluster. c) Map of *Ae. mutica* observations and their PCA cluster overlaid on Köppen-Geiger classifications (Beck *et al*. 2018): cluster 1 (Central Anatolian), cluster 2 (hot temperate), and cluster 3 (South Caucasus). Sequenced individuals are indicated by colour: T05 (light blue), T19 (dark green), T2130012 (red).

### Low-coverage HiFi sequencing and genome assembly

Over 5 million HiFi CCS reads were produced for each individual, corresponding to approximately 9× coverage per haplotype. Using these data, we produced two partially phased *de novo* assemblies for each individual using hifiasm. BUSCO assessments of these assemblies showed a high level of gene-space covered against the viridiplantae_odb12 database (822 genes; Table 1), with duplicated BUSCOs accounting for 10.7% to 12.8% of the gene set, consistent with partial rather than complete haplotype separation. Following reference-guided scaffolding to T2130012 haplotype 1, 94.4% to 96.3% of each contig-level assembly was placed onto the seven reference pseudomolecules (Table S1). The resulting scaffolded assemblies were used for identifying synteny and structural variants.

**Table 1.**
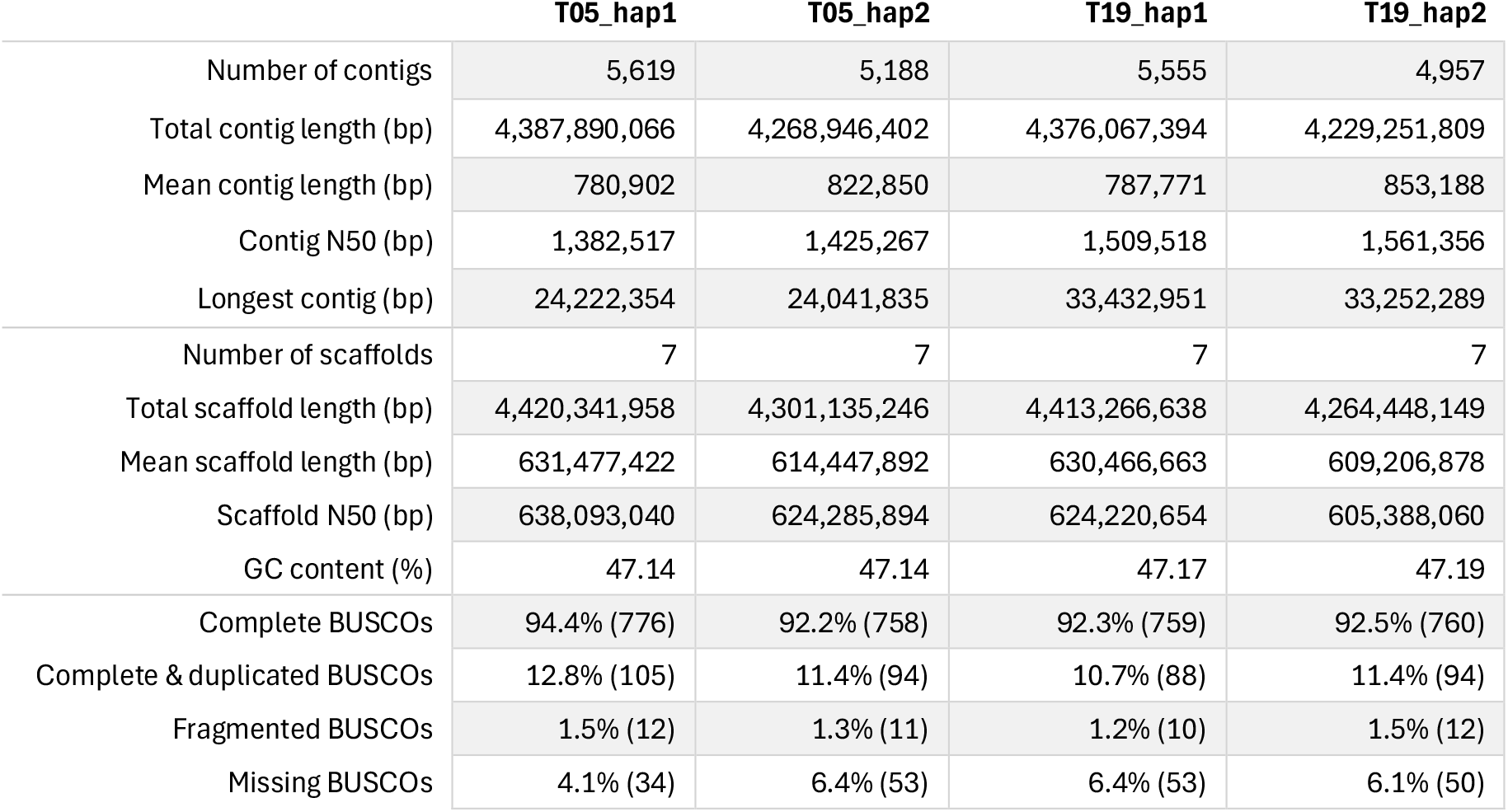
Assembly statistics for partially phased hifiasm assemblies scaffolded to T2130012 haplotype 1.

|  | T05_hap1 | T05_hap2 | T19_hap1 | T19_hap2 |
| --- | --- | --- | --- | --- |
| Number of contigs | 5,619 | 5,188 | 5,555 | 4,957 |
| Total contig length (bp) | 4,387,890,066 | 4,268,946,402 | 4,376,067,394 | 4,229,251,809 |
| Mean contig length (bp) | 780,902 | 822,850 | 787,771 | 853,188 |
| Contig N50 (bp) | 1,382,517 | 1,425,267 | 1,509,518 | 1,561,356 |
| Longest contig (bp) | 24,222,354 | 24,041,835 | 33,432,951 | 33,252,289 |
| Number of scaffolds | 7 | 7 | 7 | 7 |
| Total scaffold length (bp) | 4,420,341,958 | 4,301,135,246 | 4,413,266,638 | 4,264,448,149 |
| Mean scaffold length (bp) | 631,477,422 | 614,447,892 | 630,466,663 | 609,206,878 |
| Scaffold N50 (bp) | 638,093,040 | 624,285,894 | 624,220,654 | 605,388,060 |
| GC content (%) | 47.14 | 47.14 | 47.17 | 47.19 |
| Complete BUSCOs | 94.4% (776) | 92.2% (758) | 92.3% (759) | 92.5% (760) |
| Complete & duplicated BUSCOs | 12.8% (105) | 11.4% (94) | 10.7% (88) | 11.4% (94) |
| Fragmented BUSCOs | 1.5% (12) | 1.3% (11) | 1.2% (10) | 1.5% (12) |
| Missing BUSCOs | 4.1% (34) | 6.4% (53) | 6.4% (53) | 6.1% (50) |

### Landscape of structural variation

Using SyRI, we detected many novel structural variants (SVs) in *Aegilops mutica* compared to the reference and between partially-phased haplotypes of a single individual (Figure 3). Structural variants did not intersect any of the major domestication genes we examined, including those controlling flowering time (*Vrn1, Vrn2, Vrn3, Ppd1*), grain hardness and shattering (*Q*/q, Btr1, *Btr2*), or plant height (*Rht1*).

**Figure 3.**
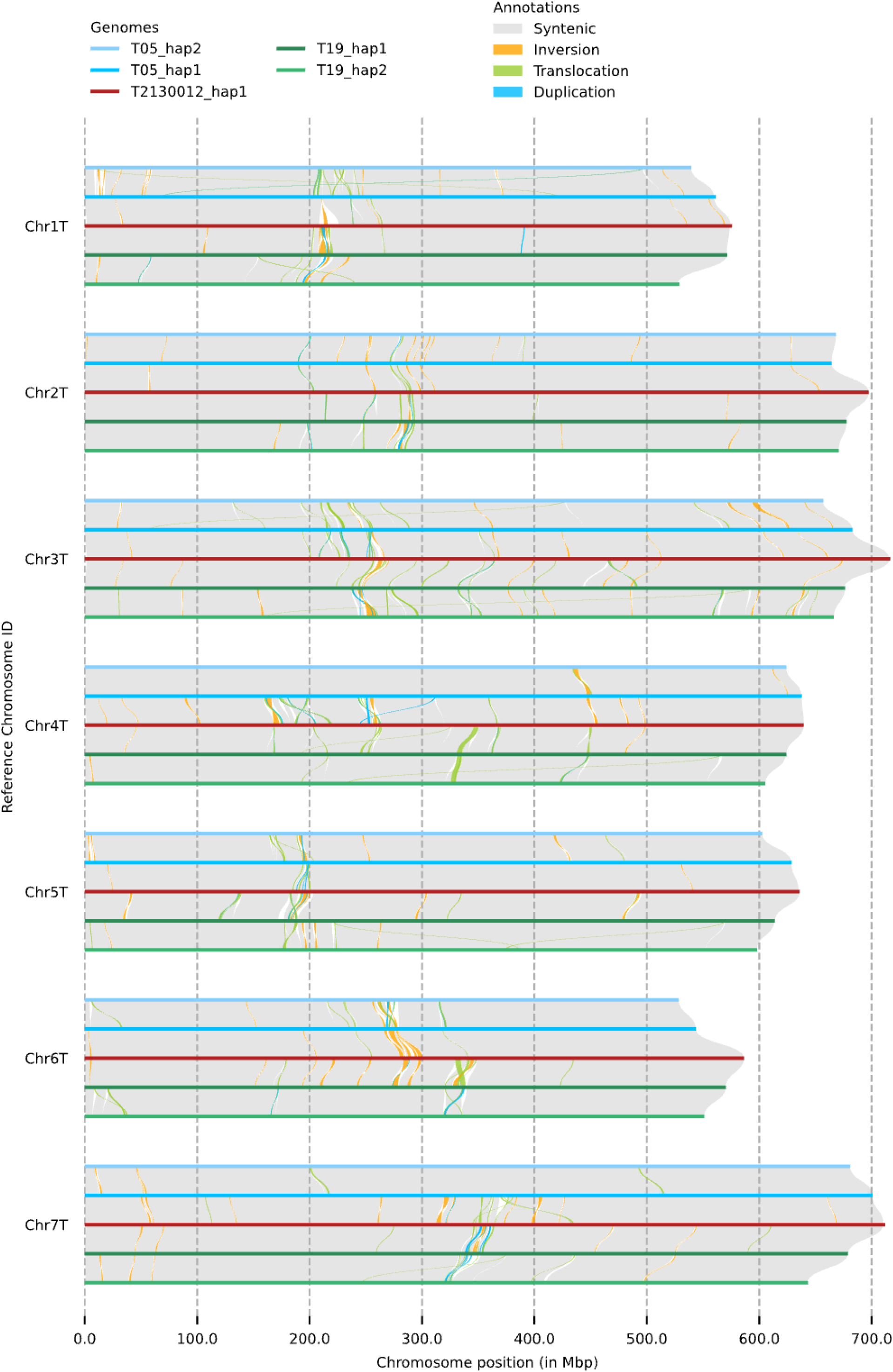
Synteny and structural rearrangements between haplotypes of *Aegilops mutica*. Synteny plot visualizing inversions, translocations, and duplications over 500 Kb between T05, T19, and the reference, T2130012.

#### Reference-based

When calling SVs using T2130012 haplotype 1 as a reference, we detected 145,871 to 155,535 SVs per haplotype. Structural variant size distribution varied by type: inversions were both the largest and the most size-variable type (mean 196 kb, median 16 kb), whereas duplications (mean 6.8 kb, median 5.7 kb) and translocations (mean 9.0 kb, median 6.4 kb) were more tightly distributed (Figure 4a). The largest SVs detected in each assembly were as follows: a 7.5 Mb inversion on chromosome 4T of T05 haplotype 1 (INV57854, Chr4T:167880167-175337548), an 8.4 Mb inversion on chromosome 6T of T05 haplotype 2 (INV56521, Chr6T:283510599-291886718; corresponding to a cluster of inversions in T05 haplotype 1, visible in Figure 3), a 4.7 Mb translocation on chromosome 6T of T19 haplotype 1 (TRANS65393, Chr6T:329617749-334324961), and a 4.6 Mb inversion on chromosome 6T of T19 haplotype 2 (INV57643, Chr6T:283660186-288282737). Averaged across the four new assemblies, the fewest SVs were detected on chromosome 4T (mean 17,573) and the most detected on chromosome 7T (mean 25,128).

**Figure 4.**
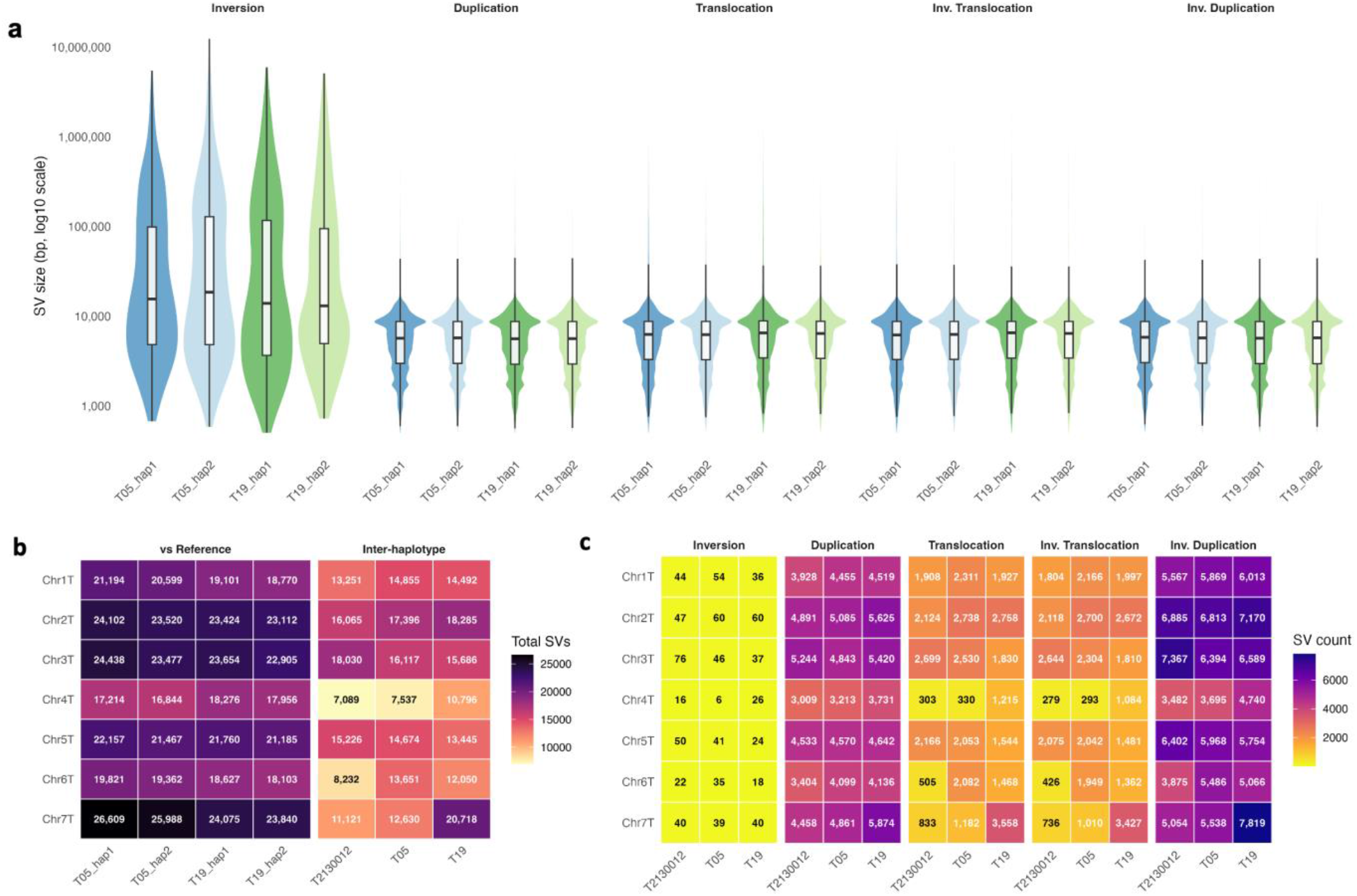
The structural variant landscape of Ae. mutica assemblies. a) Size distribution of reference-based structural variants per assembly, by type, on a log10 scale. Inversions have a varied size and tend to be larger than the other types. b) Heatmap of structural variant counts per chromosome. Reference-based SVs (left) and single individual inter-haplotype SVs (right). c) Heatmap of inter-haplotype structural variant counts per chromosome, split by type of SV. There are few inversions per chromosome but numbers of translocations and duplications are higher and vary more per chromosome.

Specifically, the number of inversions per chromosome ranged from 41-92, translocations from 2,336-4,342, and duplications – either tandem or distal – from 6,167-9,620, being fairly consistent across assemblies (Figure S2).

#### Inter-haplotype

We detected fewer SVs between the haplotypes of a single individual than between any haplotype and the reference genome, suggesting the two haplotypes of a single individual are more similar to each other than to the reference (Figure 4b). Between the two haplotypes of a single individual, 96,860-105,472 structural variants segregate. The largest SVs in T05 and T19 were a 3.1 Mb inversion on chromosome 5T and a 2.4 Mb inversion on chromosome 2T, respectively. The most SVs were detected on chromosome 2T of T05 and chromosome 7T of T19, compared to chromosome 3T of T2130012. As with the reference-based SVs, there were few inversions per chromosome (6-76), a middling number of translocations – albeit of a wider range (303-3,558), and a larger number of duplications (3,009-7,819). These counts varied more per chromosome than the reference-based SVs, with a particularly low number of translocations on chromosome 4T in T05 and T2130012 compared to T19 (Figure 4c).

### GO terms enriched in structural variants

Gene Ontology (GO) enrichment analysis of genes within or overlapping the boundary of structural variants identified overrepresentation of certain processes. A quarter of annotated reference genes which intersected a structural variant had sufficient GO terms to be included in the enrichment analysis. Among these genes, several clusters of GO terms were consistently high across assemblies, particularly in duplication SVs (Figure S3). The first cluster is associated primarily with inverted duplications and secondarily with inverted translocations and regular duplications; it contains GO terms surrounding regulation of exocytosis and secretion (mean adjusted p = 3.86×10^-11^) including syntaxin binding/SNARE binding (mean adjusted p = 3.07×10^-9^) and GTPase activity (mean adjusted p = 1.44×10^-7^). The second cluster, associated primarily with duplications followed by inverted duplications/translocations, contains regulation of ribosome biogenesis (mean adjusted p = 5.42×10^-9^), rRNA processing (mean adjusted p = 5.41×10^-9^), and RNA polymerase I transcription (mean adjusted p = 4.67×10^-8^). The final consistently enriched cluster is strongly associated with duplications and surrounds energy metabolism: ATP synthase (mean adjusted p = 3.16×10^-6^), NADH dehydrogenase (mean adjusted p = 2.07×10^-4^), and oxidative phosphorylation (mean adjusted p = 0.0068). Simple inversions and translocations are absent from background enrichment. Several clusters of GO terms were uniquely enriched within inversions on both haplotypes of each individual, these centred around stress responses and included aldehyde detoxification and unfolded protein response in T05, and herbicide detoxification and oxidative defence in T19.

### Repeat and transposable element content

Repeat content of the four new *Aegilops mutica* haploid genome assemblies ranges from 78.27% to 82.80% (Figure S4). LTR retrotransposons are the dominant class, covering approximately 55% of each assembly, of which elements of the Ty1/Copia and Ty3 superfamilies represent 17–18% and 36–37% of total assembly length respectively. Kimura distance distributions of TE subclasses were comparable between T05 and T19 and between haplotypes of the same individual (Figure S5).

## Discussion

Here, we present four new partially phased assemblies from two *Ae. mutica* individuals, filling the gap in genomic resources for this species. Crucially, these accessions were originally collected from opposite ends of the species’ range from bioclimates that are different from that of the reference genome. We have shown how common structural variations are within *Ae. mutica* and how the landscape of these SVs differs between individuals, particularly when comparing inter-haplotype SVs.

### Structural variants in climatically distinct individuals

Inversions were the SV type with the largest median and mean size. Large inversions contain numerous genes and can link alleles together in largely non-recombining haplotype blocks by suppressing recombination, these can persist and then spread through a population (Shipilina *et al*., 2022). Haploblocks like this have often been linked to ecotypes as a form of adaptation for survival in certain environments as well as acting as supergenes (Joron *et al*., 2011; Todesco *et al*., 2020; Meyer *et al*., 2024). The large inversions seen in both T05 and T19 represent candidate structural variants that warrant investigation for potential roles in local adaptation (Kim *et al*., 2022). The number of SVs detected here, between 145,871 and 155,535, is far higher than were found in *Aegilops* species from the Sitopsis clade compared to the polyploid wheat B subgenome (between 37,039 and 37,721; Li *et al*., 2022). This disparity is partially explained by difference in SV detection method; the program used by Li *et al*. 2022, Sniffles, fails to detect 20% of tandem and 40% of distal duplications compared to SyRI (Goel *et al*., 2019). The rest could be explained by differences in alignment and sequence similarity: higher overall synteny would be found between multiple assemblies of *Ae. mutica*, leading to more individual SVs being detected, whereas when comparing between different species, non-syntenic regions could instead be called simply as highly divergent regions with no SVs detected. Additionally, the SV counts are likely a conservative estimate, as reference-guided scaffolding can mask rearrangements in regions of low synteny.

The majority of the *Ae. mutica* genome has been transferred into wheat via introgression lines from three Turkish *Ae. mutica* accessions, one of them being the reference T2130012 (King *et al*., 2026), however, assemblies of T05 and T19 harbour over one hundred thousand SVs compared to the reference. These SVs are representative of genomic diversity from across the species range that has not been introgressed into wheat. Both accessions are of interest for crop development due to being bioclimatically distinct from the reference accession. The more geographically isolated accession of T19 is of particular interest for winter wheat given the colder climate from which it was collected. None of the major domestication loci were situated within a large structural variant. Rather than a null result, we regard this as an informative positive control: these loci are functionally constrained and expected to be structurally conserved across the species. Their presence in syntenic, variant-free regions in all four haplotypes indicates that the assembly and SV-calling pipeline is not generating spurious rearrangements in gene-rich, conserved sequence.

### Enrichment of GO terms associated with structural variants

Across all four haplotypes of the two individuals there is a pattern of enrichment within GO-annotated genes overlapping structural variants. Three core groups of GO terms are enriched – energy metabolism, ribosome biogenesis, and transmembrane trafficking – particularly within duplications. This may indicate a prevalence of copy number variation for the associated genes. Additionally, the uniquely enriched stress-related GO terms present within inversions on both haplotypes of an individual could be evidence of inversion haploblocks maintaining co-adapted genes by suppressing recombination, however, no conclusions can be drawn between these samples and local adaptation due to only a single individual representing each environment. Further population level analysis would be required to ascertain whether any inversions are correlated with a particular environment.

### Repeat content and transposable elements

The repeat content of the four new assemblies is slightly higher than the 77.01% reported for the reference (Grewal *et al*., 2025), but is broadly comparable to other Triticeae species (Feldman and Levy, 2023) such as 75.2% in *Aegilops speltoides* (Avni *et al*., 2022) and 84.4% in *Aegilops tauschii* (Luo *et al*., 2017). LTR retrotransposon coverage is marginally lower than the 59% found in the reference genome, yet the Ty1/*Copia* and Ty3 proportions fall within 1% of reference values, indicating that the repeat landscape is stable across the species range despite the structural variation described above. The assembly-wide proportion of LTR classes Ty1/*Copia* and Ty3 LTRs in all four assemblies falls within 1% of those measured in the reference genome: 17-18% and 36-37%, respectively. These two classes of LTR often underpin insertions and recombination-mediated rearrangements in plants, particularly through biotic and abiotic stress-activated retrotransposition bursts (Papolu *et al*., 2022); this is reinforced here by the high numbers of both SVs and Ty1 and Ty3 LTRs.

### Conclusions

We generated four new partially phased *Ae. mutica* assemblies and identified extensive structural variation both among individuals and between haplotypes. This study highlights the benefits and limitations of using a reference-guided assembly. Our approach of forgoing optical mapping and choosing low-depth long-read sequencing reduces the cost of investigating the prevalence of structural variants. However, this necessitated scaffolding the assemblies to a reference genome, which inevitably constrains the detection of novel rearrangements. Despite this limitation, we detected many SVs, both between the haploid genomes of an individual and when compared to the reference genome. This highlights the complexity of large plant genomes and shows that a single reference genome is insufficient to fully describe a complex, diverse species such as *Ae. mutica* (Edwards *et al*., 2025). These assemblies provide a foundational step toward a species-level pangenome, enabling the development of structural variant markers for beneficial traits.

Future work should prioritise linking these structural variants to phenotypic data in order to advance climate-resilient wheat breeding. Similarly, future conservation strategies must account for these distinct populations to fully represent the species’ biodiversity and capture its adaptive potential. However, drawing from genebank accessions may be the only way to do this, as the Armenian population at Erebuni reserve was not seen between 2021 and 2023 (Nersesyan, 2024). This endangered and underutilised species should continue to be researched for potential agricultural benefits, and key populations should be closely monitored as a species of conservation priority.

## Abbreviations

BUSCO: Benchmarking Universal Single Copy Orthologs
CCS: circular consensus sequencing
CWR: crop wild relative
GO: gene ontology
LTR: long terminal repeat
PCA: principal component analysis
SNP: single nucleotide polymorphism
SSR: simple sequence repeat
SV: structural variant
TE: transposable element

## Supplemental Material statement

Supplemental material for this article:

- Table S1: Statistics for the initial genome assemblies before scaffolding.
- Figure S1: BUSCO completeness scores used to select the best assembly settings.
- Figure S2: Referenced-based structural variant counts per chromosome.
- Figure S3: GO term enrichment for different types of structural variants.
- Figure S4: Repetitive sequence content in the new genome assemblies.
- Figure S5: Estimated age and diversity of repetitive sequences.

## AI Declaration statement

Claude Sonnet 4.6 and GPT-5.2 (accessed May 2026) were used in the creation of scripts for TE, GO term enrichment, and structural variant analysis, however, all code was manually reviewed, tested, and revised by the authors.

## Data Availability statement

The genome assemblies and raw sequencing reads generated in this study have been deposited in ENA under project accession PRJEB126962/ERP206544. Structural variant calls as SyRI outputs and VCFs are available from Figshare (DOI: 10.6084/m9.figshare.33841576).

## Author Contributions

**Conceptualisation:** ALT, RS. **Methodology:** ADT, ALT, MH, RS. **Formal Analysis:** ALT. **Investigation:** ALT. **Resources:** ADT, ALT, MH, RS, WP. **Writing – Original Draft:** ALT. **Writing – Review & Editing:** ADT, ALT, BK, GB, MH, RS, WP. **Visualisation:** ALT. **Supervision:** ADT, GB, MH, RS (lead).

## Acknowledgments

We would like to thank the following: Dr Hakan Özkan from Çukurova University for information on *Ae. mutica* populations in Central Türkiye; Dr. Anush Nersesyan from Armenian National Academy of Sciences and Dr. Margarita Harutyunyan from Armenian National Agrarian University for information on *Ae. mutica* populations in Armenia; Julie Fortune, G Cuthill, and the technicians at SRUC for care of plants in the glasshouse.

This work was supported by Scotland’s Rural College (SRUC PhD studentship 1036656). MH is supported by a NERC Independent Research Fellowship (NE/R015686/1) and a UKRI Frontier Research Guarantee Grant (EP/X027570/1). Sequencing was carried out by Edinburgh Genomics, The University of Edinburgh, which is partly supported with core funding from the Natural Environment Research Council (UKSBS PR18037). We acknowledge Research Computing at the James Hutton Institute for providing computational resources and technical support for the “UK’s Crop Diversity Bioinformatics HPC” (BBSRC grants BB/S019669/1 and BB/X019683/1), use of which has contributed to the results reported within this paper.

