## Supplemental material for "Genome assemblies of climatically distinct *Aegilops mutica* reveal extensive structural variation"

Ashley L. Tuffin, Georgios Banos, Benjamin Kilian, Wayne Powell, Alex D. Twyford, Matthew Hartfield, and Rajiv Sharma.

|  | T05_hap1 | T05_hap2 | T19_hap1 | T19_hap2 |
| --- | --- | --- | --- | --- |
| Number of contigs | 7,661 | 6,600 | 7,900 | 6,041 |
| Total contig length (bp) | 4,646,364,162 | 4,434,055,336 | 4,635,587,486 | 4,435,888,234 |
| Mean contig length (bp) | 606,496 | 731,692 | 586,783 | 734,297 |
| Contig N50 (bp) | 1,278,510 | 1,364,070 | 1,421,823 | 1,500,084 |
| Unplaced length | 258,474,096 | 165,108,934 | 259,520,092 | 206,636,425 |
| % anchored to Chr1T-7T | 94.44 | 96.28 | 94.40 | 95.34 |

### Table S1 Metrics of unscaffolded HiFiasm assemblies

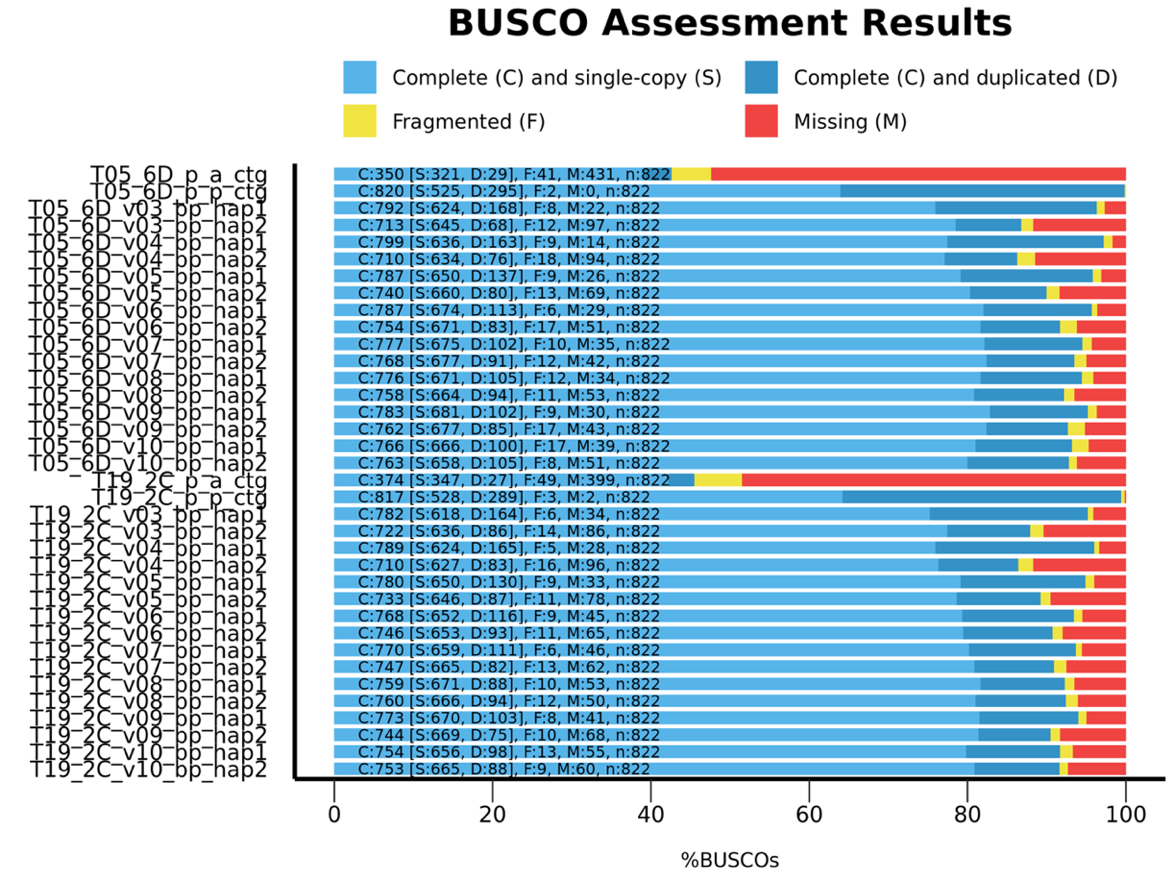

### Figure S1 BUSCO assessment results for HiFiasm similarity threshold (s) values 0.55 (v3) to 0.05 (v10) and primary/alternative assemblies.

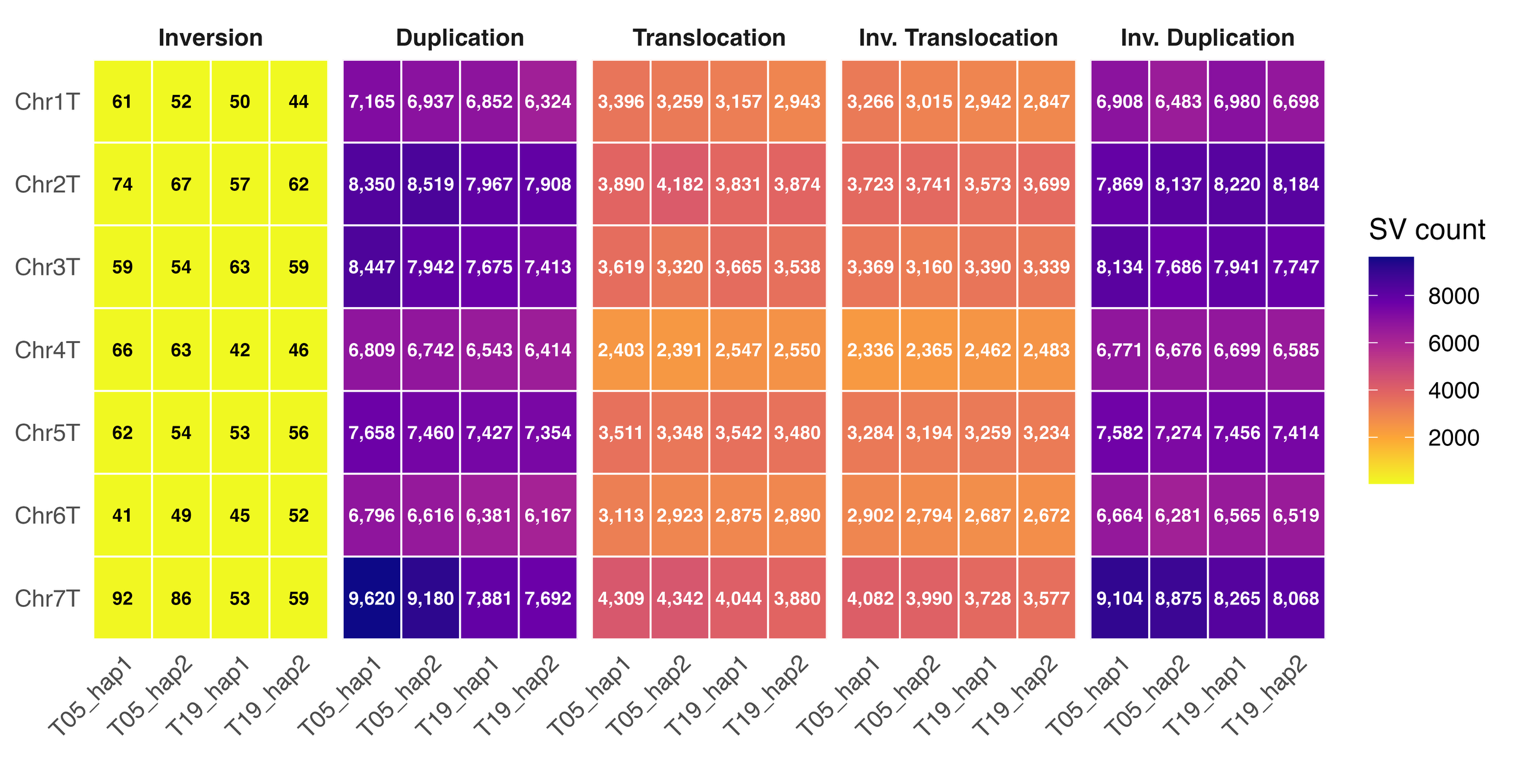

### Figure S2 Structural variant type counts per chromosome per assembly, detected against the reference genome of T2130012 haplotype 1.

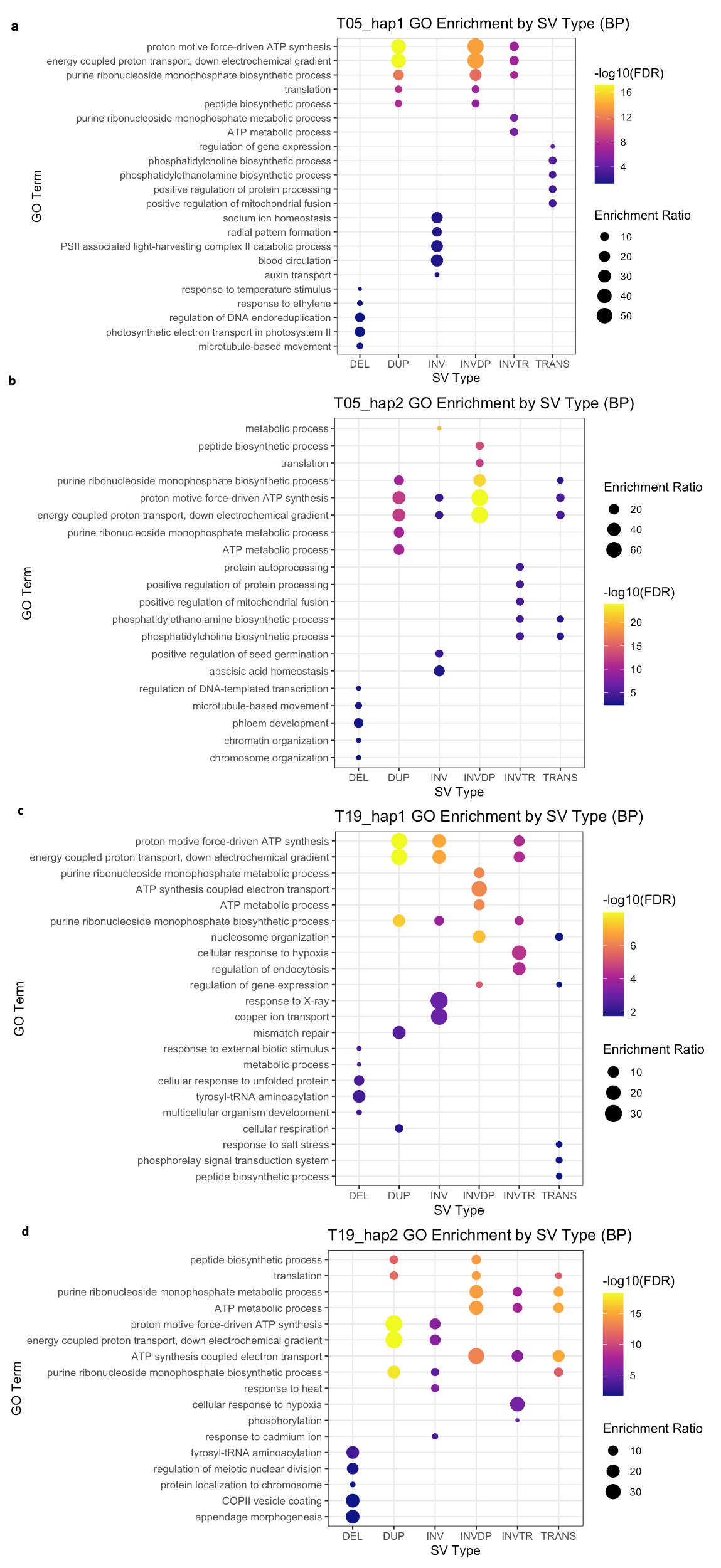

### Figure S3 GO term enrichment plots for biological processes (BP) in T05 and T19 assemblies in different structural variant types.

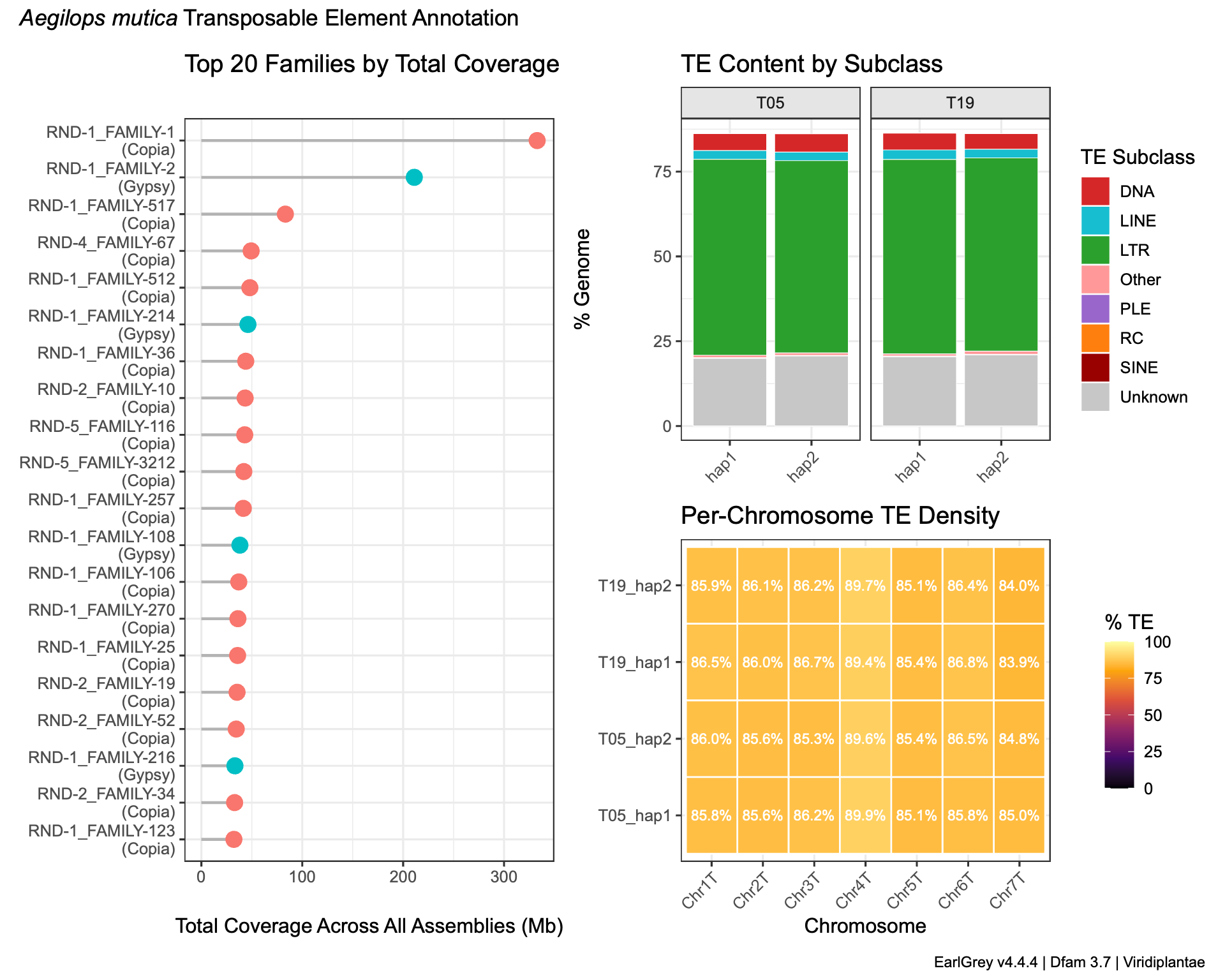

### Figure S4 Quantity of transposable elements within T05 and T19 assemblies.

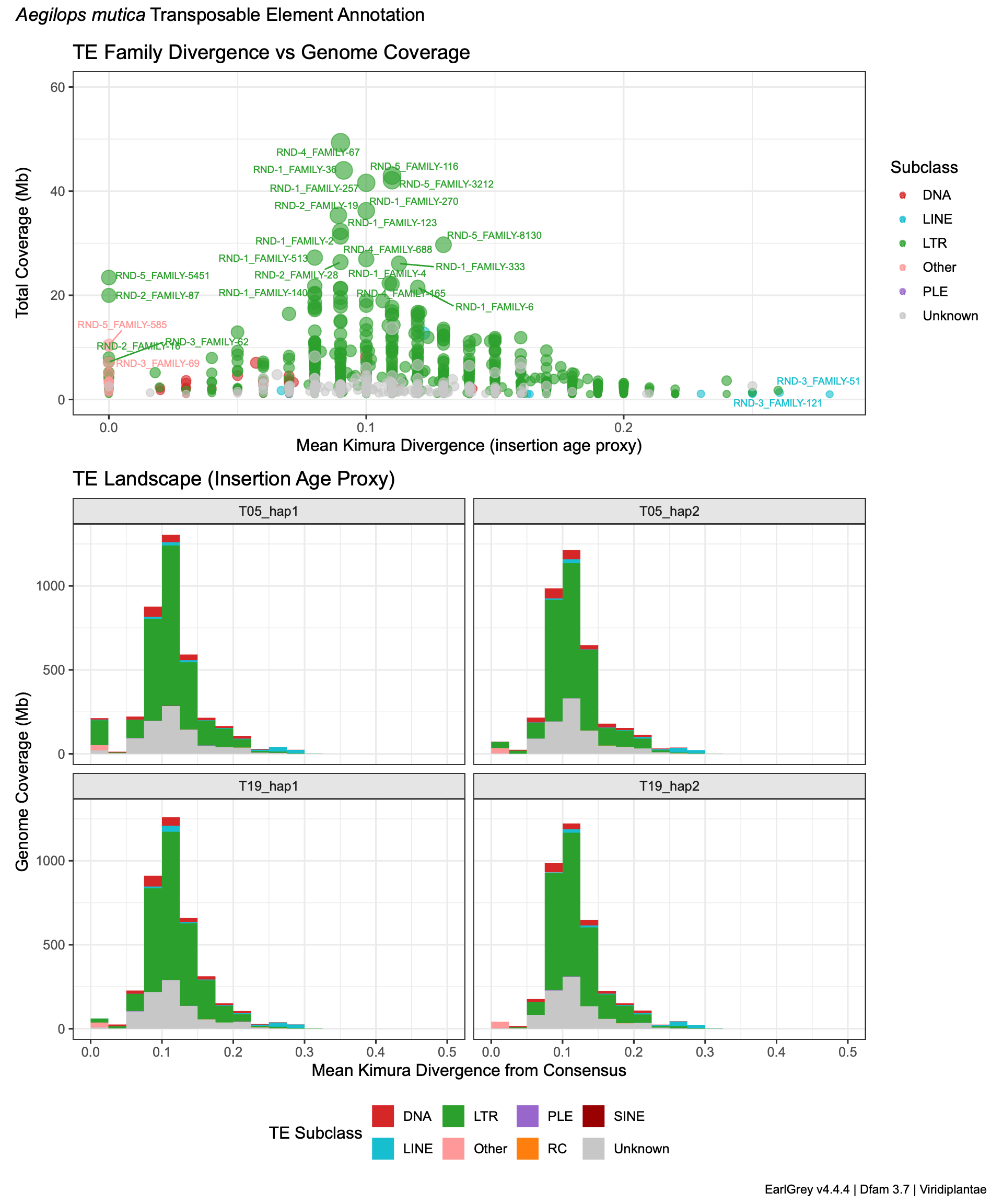

### Figure S5 Kimura divergence of transposable elements subclasses within T05 and T19 assemblies.
